# Winter rye root growth and plasticity in response to nitrogen and phosphorus omission under field conditions

**DOI:** 10.1101/2025.01.22.634240

**Authors:** John Kormla Nyameasem, Sabine J. Seidel, Sofia Hadir, Gina Lopez, Sara L. Bauke, Ixchel M. Hernandez-Ochoa

## Abstract

Nitrogen (N) and phosphorus (P) deficiencies can significantly reduce crop yield. Despite their importance, the impacts of N and P deficiencies under field conditions on cereal roots, particularly winter rye, remain poorly understood. This study investigates the effects of N and P deficiencies on winter rye growth and root architecture under field conditions. A sampling campaign was conducted during the 2022 season at the long-term fertilizer experiment Dikopshof, Germany. Four fertilizer treatments were chosen: (1) fully fertilized with manure (NPKCa+m+s), (2) fully fertilized without manure (NPKCa), (3) N omitted (_PKCa), and (4) P omitted (N_KCa). Shoot biomass was assessed at five growth stages, alongside with topsoil root biomass, number of nodal roots and tillers, and root angle. The results showed that shoot and root biomass were highest in the NPKCa+m+s treatment and lowest under N omission. Although the treatment ranking of root traits varied between dates, a trend for an enhanced number of roots in the N and P omission treatments was observed around flowering. P omission fostered an increased number of tillers and N omission caused steeper root angles compared to other treatments. These findings demonstrate the strong impact of the environment and development stage on root phenotypic plasticity.

## 2) Introduction

Nitrogen (N) and phosphorus (P) are critical nutrients that are central to crop productivity, significantly influencing plant growth, development, and yield. N is key for amino acid and protein synthesis, while P is essential for energy transfer, photosynthesis, and structural development (Uga et al., 2015). However, the availability of these nutrients is often limited in agricultural soils, constraining crop yields, and quality (Hammer et al., 2009). Addressing these constrains requires improving crop nutrient use efficiency to tackle the twin challenges of global food security and environmental sustainability. Among the various strategies, enhancing the functional plasticity of root systems emerges as a key approach to improve nutrient acquisition and crop performance under resource-limited conditions.

Roots play a vital role in plant adaptation to environmental and edaphic stresses. The architecture of the root system—comprising traits such as root biomass, length, and angle—is key to exploring soil and acquiring nutrients (Lynch, 1995; Lynch, 2019). Environmental factors, including soil water availability, salinity, and nutrient distribution, further influence root growth dynamics, modulating root system depth, angle, and efficiency in nutrient and water uptake (Manschadi et al., 2008; Oyanagi et al., 1993). Recent studies highlight strategic adjustments in root growth patterns that enable plants to optimize resource use efficiency in suboptimal environments (Lynch, 2019). Among these traits, root angle is a fundamental determinant of root placement within the soil profile. Steeper root angles promote deeper soil penetration, enhancing access to mobile resources like nitrate and water under drought or low-N conditions. For instance, simulations have demonstrated an 11% increase in N uptake and a 4% biomass gain under low-N conditions in genotypes with steeper root angles (Schneider et al., 2022; Trachsel et al., 2013). Conversely, shallower root angles facilitate topsoil exploration, improving the acquisition of immobile nutrients such as phosphorus (Bonser et al., 1996; Liao et al., 2001; Lynch & Brown, 2001). These contrasting strategies underscore the importance of root angle plasticity in mediating crop responses to nutrient constraints and ensuring resilience under varying soil conditions (Lynch, 2018).

Cereal crops, which form the foundation of global food systems, exhibit distinct root system responses to N and P deficiencies. A recent review reported that under N deficiency, crop root length and biomass decreased by 9% and 7%, respectively, but root length per shoot biomass increased by 33%, alongside a 44% enhancement in the root-to-shoot ratio, reflecting carbon allocation strategies for nutrient foraging (Lopez et al., 2023). On the other hand, P deficiency induced more substantial reductions in root length (14%) and biomass (25%) but increased root length per shoot biomass by 51%, though the root-to-shoot ratio exhibited inconsistent trends (Lopez et al., 2023). These responses highlight trade-offs between growth and nutrient acquisition, with enhanced topsoil root proliferation compensating for P scarcity, while N limitation restricts overall biomass due to carbon allocation constraints (Bonser et al., 1996; Mao et al., 2018).

Long-term experiments serve as an important platform for research, however, studies mostly focus on the above-ground traits, leaving the effects of adaptation on the root architecture to fertilizer omission largely unknown (Siddiqui et al., 2021). Given the potential of low-input agriculture for sustainable crop production, further investigation is essential. Despite extensive research on the effects of N and P availability on shoot performance, the specific impacts of nutrient omissions on root system architecture under field conditions, particularly in cereals such as winter rye (*Secale cereale* L.), remain underexplored (Lopez et al., 2023). Renowned for its adaptability to nutrient-poor soils and challenging growing environments, winter rye serves as an excellent model for investigating root-shoot interactions under nutrient stress (Arsova et al., 2020). Its capacity to thrive in suboptimal conditions offers a unique opportunity to study how root angle and associated traits respond to varying nutrient availability during the growth period. This study aims to address this knowledge gap by examining the effects of nutrient omissions on root and shoot traits in winter rye. By examining morphological root trait adaptations that enhance resource use efficiency, the study seeks to provide valuable insights for developing crop management and breeding strategies to improve productivity and resilience in nutrient-limited environments.

## 3) Method

### Experimental design

A sampling campaign was conducted in 2022 at the long-term fertilizer experiment (LFTE) Dikopshof established in 1904 near Cologne, Germany. The general soil type is classified as a Haplic Luvisol derived from loess above sand with a silty loam (topsoil) and (silty) clay loam (below 30 cm soil depth). The five-year crop rotation at the LTFE Dikopshof comprises sugar beet (*Beta vulgaris*), winter wheat (*Triticum aestivum* L.), winter rye, Persian clover (*Trifolium resupinatum* L.), and potato (*Solanum tuberosum* L.). The experiment is a non-randomized block design without replicates and comprises seven treatments: NPKCa+m+s (+m stands for farmyard manure fertilization and +s stands for supplemental mineral fertilization), NPKCa, _PKCa, N_KCa, NP_Ca, NPK_, and no fertilizer (_ stands for the omission of the corresponding nutrient, Ca stands for lime). After harvesting the preceding crop, cattle farmyard manure is supplied on sugar beet, potato, and winter rye plots at a rate of 60 t ha^-1^ per five-year rotation (fresh matter, treatments “+m”). The fertilization management has not changed since 1953, except for a slight increase of the N fertilizer application (+ 30 kg N ha^-1^) on winter wheat in some treatments, which occurred in the 1980s. More details are presented in Seidel et al. (2021).

### Sampling and analysis

Four treatments were considered in this study: Fully fertilized plus manure (treatment NPKCa+m+s), fully fertilized with mineral fertilizer only (NPKCa), N omission (_PKCa), and P omission (N_KCa). Winter rye shoot biomass was estimated destructively on five dates (16/03, 04/04, 29/04, 27/05, and 21/06) from tillering to flowering by cutting four times 50 cm of a row. Simultaneously, root coring using an auger (9 cm inner diameter) was conducted to sample the roots in the regularly ploughed topsoil (0-30 cm). Roots per auger were then carefully cleaned with tap water, and nodal root angles, the number of tillers, and the number of nodal roots emerging from shoot tissue (root number) were estimated manually for all plants. The angular spread of the roots was defined as the deviation angle of the two most horizontally distant shoot roots (180° would be roots at soil surface, Figure S1). The samples were then sieved (2 mm and 0.63 mm) and sorted to remove the debris from the samples. Roots were then dried weighted to derive dry matter root biomass and determine C and N tissue content.

Four soil samples per treatment and sampling date were taken from the ploughed topsoil (0-30cm) with a Pürkhauer auger. The samples per treatment were then pooled together and frozen. After thawing, the soil was analyzed for mineral nitrogen content (N_min_) by extraction with potassium sulfate solution. The N_min_ concentrations in the extracts were measured by a Skalar Continuous Flow Analyser (Skalar Analytical B.V., Breda, Netherlands). Plant-available phosphorus (P_cal_) and potassium (K_cal_) were determined using a calcium acetate calcium lactate extract as described by Schüller (1969). P concentration in the extracts was determined colorimetrically following molybdenum blue reaction (Murphy & Riley, 1962) on a spectrophotometer (Specord 205, Analytik Jena, Germany). Volumetric soil water content (VWC) at 3 cm, 30 cm and 60 cm soil depth was measured using the FDR moisture sensor HH2 within ML3 Theta Probe (ecoTech Umwelt-Meßsysteme GmbH, Bonn, Germany) at winter rye flowering on 27/05/2022.

### Management

After farmyard manure application on 31/10/2021 and ploughing on 07/11/2021, winter rye was sown on 09/11/2021. Mineral P and N were applied on 28/03/2022 and the second N-fertilization (only NPKCa+m+s) was applied on 10/05/2022 (around BBCH 55). Harvest was on 27/07/2022.

### Statistics

The data obtained were analyzed using R software (version 4.1.1). Although the replicates are not true replicates due to the old experimental set up, a mean comparison test was carried out for all measurements, using the sub samples collected at each plot as pseudo-replicates. Data normality was tested by sampling date by using the Shapiro wilk test. The log10 or negative square root transformation data transformation was used when data was non-normally distributed. For the normally distributed data, a one-way analysis of variance (ANOVA) was conducted. Multiple comparisons between treatments were performed using Tukey’s test, and means with the same letter are considered not significantly different (Tukey test, P > 0.05). When data was not normally distributed, and data transformation was unsuccessful, an Aligned rank transform factorial ANOVA for non-parametric data was used by implementing the ARTool package (version 0.11.1). Finally, the ggcorr function in R was used to visualize the correlation coefficients between the shoot and root variables in a correlation matrix.

## 4) Results

Overall, N and P omission treatments significantly affected root and shoot biomass as well as root traits during the season. For the shoot biomass, treatment differences became more apparent during the last sampling dates, similarly for root biomass. Fertilizer treatments significantly affected root morphological traits, with greater variation observed among the different treatments. Roots number and root angle tended to increase during the season, while tiller numbers were similar across sampling dates, but with large variability among treatments.

### Soil conditions

The NPKCa+m+s resulted in the highest soil K_cal_, P_cal_ and N_min_ contents during all sampling dates (Table 1). Soil P_cal_ and K_cal_ content were the lowest in the N_KCa treatment, with more drastic differences in the soil P_cal_ content. As expected, the soil N_min_ was the highest in the NPKCa+m+s treatment with the highest values observed in the first (29.14 mg kg^-1^) and last (27.61 mg kg^-1^) sampling dates (Table 1). The rest of treatments showed considerably lower N_min_ values as well as a tendency to decrease as the season progressed. The soil N_min_ values were lowest for _PKCa in all dates (Table 1).

**Table 1.** P_cal_, K_cal_ and N_min_ soil content for four sampling dates during the 2022 growing season for winter rye.

| Sampling date | Treatment <sup>1</sup> | Soil K<sub>cal</sub> <sup>2</sup><br>(mg kg<sup>-1</sup> of soil) | Soil P<sub>cal</sub> <sup>2</sup><br>(mg P kg<sup>-1</sup> of soil) | Soil N<sub>min</sub> <sup>3</sup><br>(mg kg<sup>-1</sup> of soil) |
| --- | --- | --- | --- | --- |
| 07/04/2022 | NPKCa+m+s | 312.17 | 168.70 | 29.14 |
|  | NPKCa | 157.37 | 78.49 | 12.94 |
|  | N_KCa | 136.92 | 16.14 | 14.04 |
|  | _PKCa | 155.62 | 83.88 | 1.63 |
| 29/04/2022 | NPKCa+m+s | 287.69 | 208.90 | 5.24 |
|  | NPKCa | 135.82 | 95.84 | 8.84 |
|  | N_KCa | 103.37 | 15.59 | 2.43 |
|  | _PKCa | 160.02 | 86.22 | 1.07 |
| 27/05/2022 | NPKCa+m+s | 337.07 | 192.30 | 13.00 |
|  | NPKCa | 138.51 | 78.69 | 2.22 |
|  | N_KCa | 109.96 | 14.84 | 3.12 |
|  | _PKCa | 159.86 | 73.84 | 1.80 |
| 21/06/2022 | NPKCa+m+s | 266.64 | 198.15 | 27.61 |
|  | NPKCa | 132.31 | 87.14 | 1.90 |
|  | N_KCa | 113.21 | 15.10 | 1.20 |
|  | _PKCa | 138.06 | 71.42 | 1.18 |

<sup>1</sup>Treatment: Fully fertilized plus manure (NPKCa+m+s), fully fertilized with mineral fertilizer only (NPKCa), N omission (\_PKCa) and P omission (N\_KCa); <sup>2</sup> Soil P and K content measured using the calcium acetate-lactate extract method; <sup>3</sup> Soil mineral nitrogen content (N<sub>min</sub>) measured by extraction with potassium sulfate solution.

For soil water content, the VWC data collected around flowering on 27/05/2022, showed non-significant differences at the top 3cm (Figure S1). However, at 30 cm, the N omission treatment resulted in the highest soil moisture at ∼20%, followed by the P omission and NPKCa treatment. At 60 cm, most treatments resulted in non-significant differences, except for NPKCa+m+s, which showed the strongest decrease in soil moisture, suggesting more water uptake in the top layers, which reduced the supply to the deeper layer (Figure S1).

### Root and shoot biomass

Fertilizer treatments significantly affected root and shoot biomass over the season (Figure 1). The shoot biomass was significantly higher in the NPKCa+m+s treatment during all the sampling dates, with values ranging from 32.94 g m^-2^ at sampling date 1 to 2,123 g m^-2^ at sampling date 5 (Figure 1a). However, the N omission led to the lowest shoot biomass among all treatments during the last two dates. In the last sampling date, differences became less apparent with even the NPKCa treatment showing similar shoot biomass to the NPKCa+m+s, while N_KCa and _PKCa treatments showed the lowest shoot biomass with non-significant differences between them.

**Figure 1.**
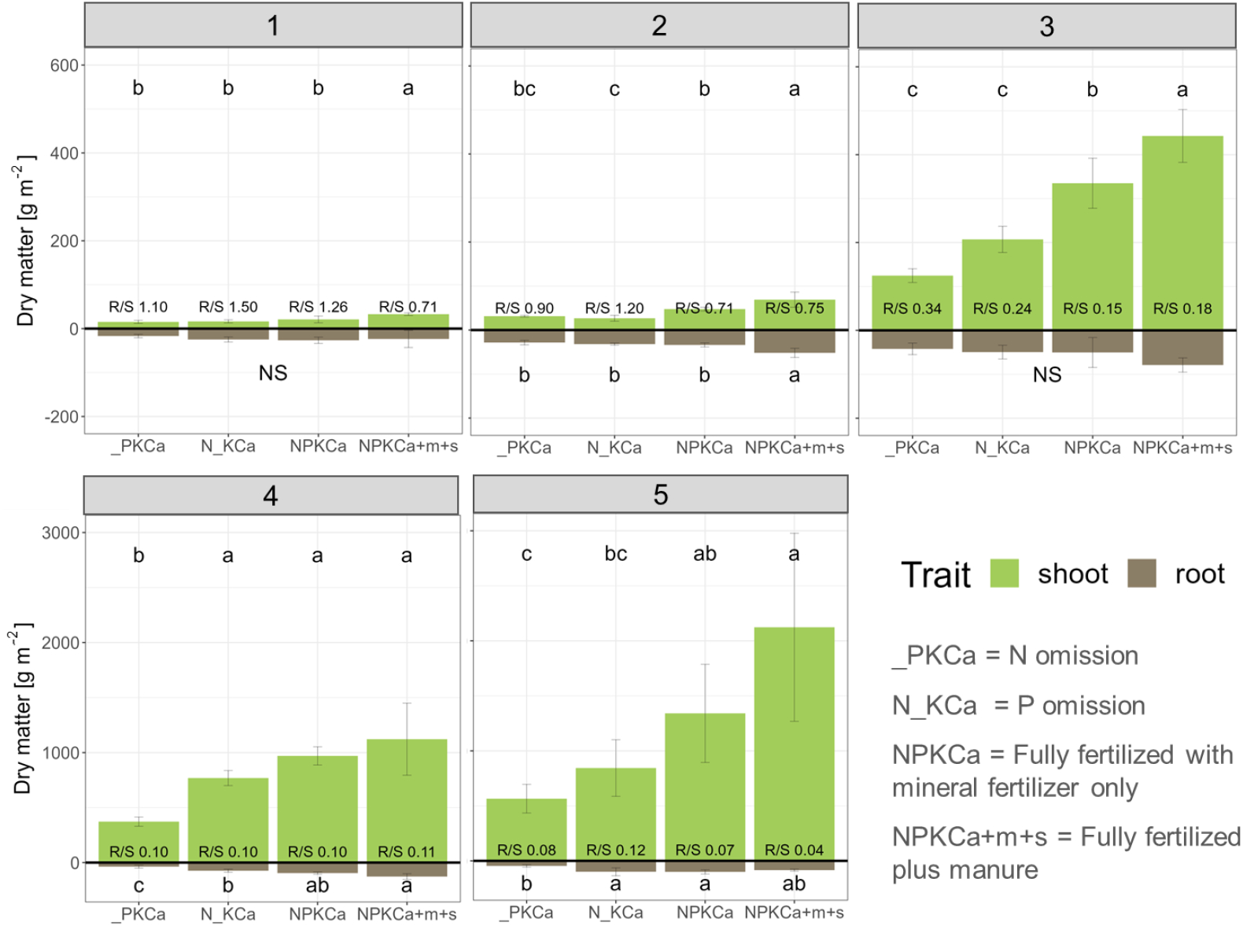
Dry matter shoot biomass (a) and root biomass in the topsoil 30 cm layer (b) in g m^-2^ and root:shoot ratio (R/S) as affected by nitrogen and phosphorus treatments over five sampling dates (1: 16/03/2022, 2: 04/04/2022, 3: 29/04/2022, 4: 27/05/2022 and 5: 06/21/2022). Treatments: Fully fertilized plus manure (NPKCa+m+s), fully fertilized with mineral fertilizer only (NPKCa), N omission (_PKCa) and P omission (N_KCa). Values followed by the same letter do not differ according to Tukey high significant difference at 5% confidence level. NS= not significant at 5% confidence level. Error bars refer to the standard deviation.

Fertilizer treatments also affected root biomass in different magnitudes, over all the sampling dates (Figure 1b). Root biomass at the first and the third sampling dates showed non-significant differences, while in the other sampling dates, the NPKCa+m+s treatment led to the highest root biomass in sampling date 2, but differences became less apparent in sampling date 4, with both fully fertilized treatments showing the highest root biomass, while the N omission treatment led to the lowest root biomass (Figure 1b). For sampling date 5, the NPKCa+m+s treatment showed a decline in root biomass, which led to non-significant difference in comparison to the N omission treatment. The P omission treatment also, caused a reduction in root biomass growth with significant differences in some sampling dates, but less than N omission. The root to shoot ratio (R/S) was highest for the P omission treatment (dates 1-3) but similar at flowering (Figure 1b).

### Morphological root traits and tillers

The measured root traits were affected by the fertilizer treatments and sampling dates, with treatments showing large variations during the season (Figure 2). For instance, the number of nodal roots was highest for NPKCa+m+s in dates 2 and 3 and lowest in dates 4 and 5, although the treatment differences were not significant (Figure 2a). The number of nodal roots did not differ significantly among the treatments in any of the dates. However, there was a trend for higher number of nodal roots in the P omission treatment compared to the fully fertilized ones in the sampling dates 4 and 5 (Figure 2a).

**Figure 2.**
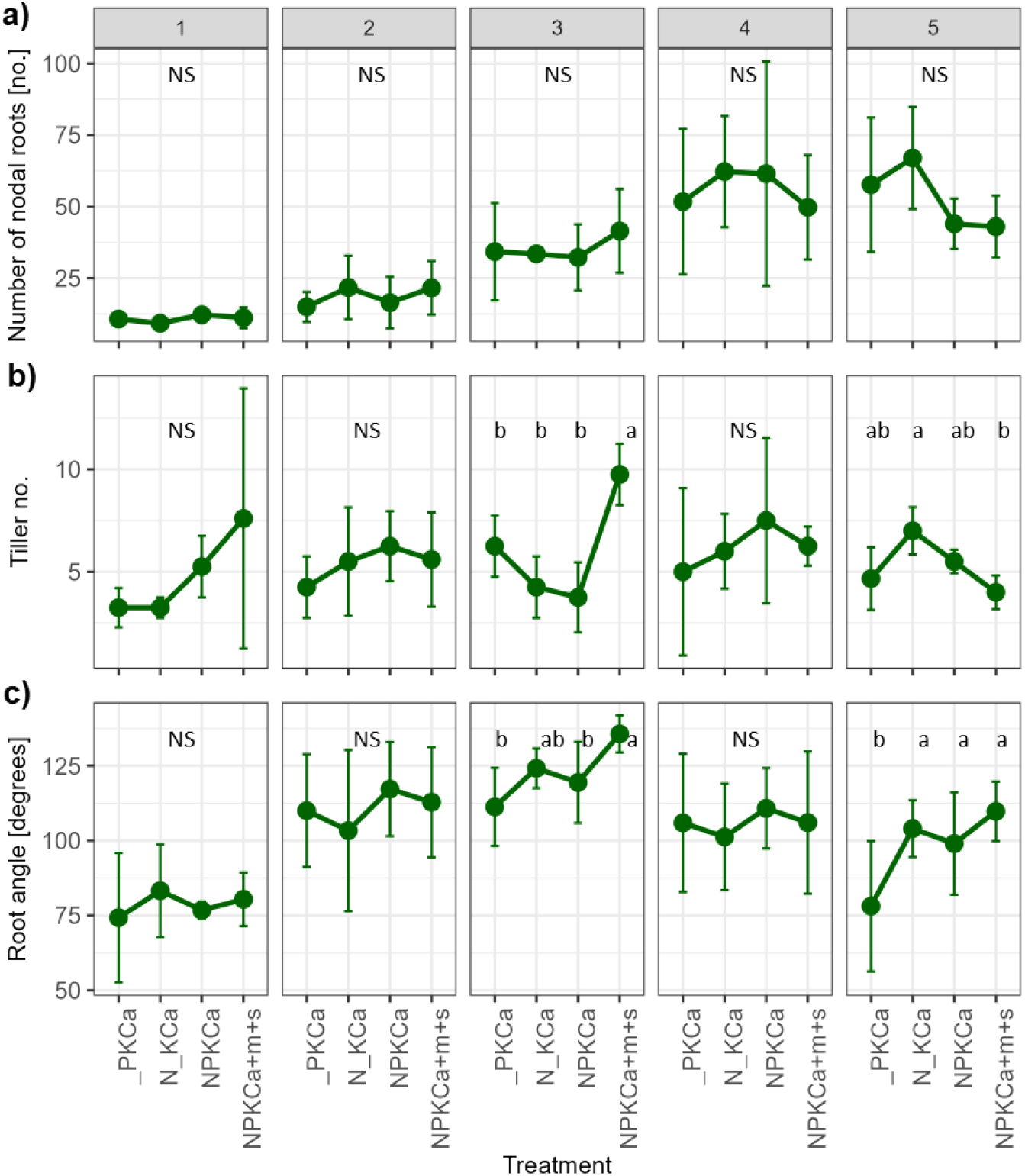
Numbers (no.) of nodal roots (a) and tillers (b) and root angle (c) for soil samples collected over five sampling dates (1: 16/03/2022, 2: 04/04/2022, 3: 29/04/2022,4: 27/05/2022 and 21/06/2022) from tillering to flowering in the top 30 cm soil layer as affected by nitrogen and phosphorus treatments. Treatments: Fully fertilized plus manure (NPKCa+m+s), fully fertilized with mineral fertilizer only (NPKCa), N omission (_PKCa) and P omission (N_KCa). Values followed by the same letter do not differ according to Tukey high significant difference at 5% confidence level. NS= not significant at 5% confidence level for the ANOVA or the Aligned rank transform for non-parametric factorial ANOVA (implemented for tillers per shoot on sampling dates 3, 4 and 5). Error bars refer to the standard deviation.

**Figure 3.**
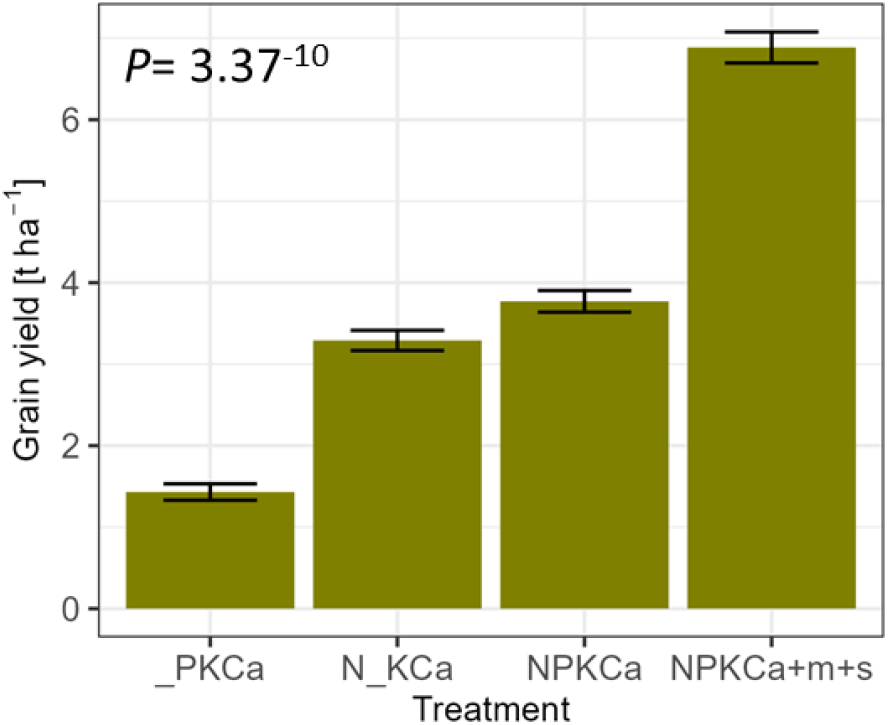
Winter rye grain yield as affected by nitrogen and phosphorus omission treatments during the 2022 season. Treatments: Fully fertilized plus manure (NPKCa+m+s), fully fertilized with mineral fertilizer only (NPKCa), N omission (_PKCa) and P omission (N_KCa). Values followed by the same letter do not differ according to Tukey high significant difference at 5% confidence level. Error bars refer to the standard deviation.

As for the tiller number, significant differences were observed during sampling dates 3 and 5, but not for the rest of sampling dates (Figure 2b). Root angles increased from values lower than 90° at the beginning of the growth period (date 1) to the largest value of 122°(mean over all treatments), observed at date 3 (Figure 2c). The largest increase of 43% was observed in the NPKCa and N_KCa treatment, and the lowest increase in the _PKCa treatment. A decrease in root angles in all treatments was observed in dates 4 and 5. The comparison of mean root angles over all treatments revealed steeper root angles in the _PKCa treatment with significant differences compared to the other treatments in dates 3 and 5. This was not the case for the N_KCa where the values were similar to the fully fertilized treatments (Figure 2c).

In general, a trend for an enhanced number of nodal roots in the N and P omission treatments was observed around flowering. Besides, P omission fostered an increased number of tillers and N omission more narrow root angles.

### C and N tissue content

Fertilizer treatments affected root and shoot C and N contents and C:N ratio. Treatment effects on shoot C content were similar among treatments and across sampling dates (Figure S3 and S4). The shoot C content was slightly higher (44.00% in average) as compared to the root C content (39.05% in average) and slightly increasing as the season progressed (Figure S3). By contrast, N content in shoots decreased from 2.8% in date 1 to 0.8% at the end of the season, similarly for roots, decreasing from 2.8% at sampling date 1, to a decrease of 76% towards the end of the season (Figure S4). Mean root N content was highest for treatment _PKCa (0.83 %) but as the total root biomass was reduced, the root N uptake (root biomass times root N content) was lowest compared to the other treatments in all dates. The C:N ratio was the lowest in shoots during sampling dates 1 to 3, but significantly increased during the last two sampling dates caused by a reduction of shoot N content (Figure S4 and S5). The NPKCa+m+s showed the lowest shoot C:N ratio in date 4. This trend was maintained during the last sampling date, though not significant due to high variation in treatment responses (Figure S5). Root C:N ratio also increased as the season progressed, though no significant differences were observed among treatments, except for the last date, where the _PKCa treatment showed a lower C:N ratio compared to the N_KCa, but the N_KCa was not significantly different to the NPKCa treatments (Figure S5).

### Grain yield

Significant differences were observed among all treatments, with the NPKCa+m+s treatment showing the highest yield with 6.8 t ha^-1^, even the NPKCa resulted in considerably lower yield of 3.8 t ha^-1^. The P and N omission treatments resulted in the lowest yields, with 53% and 80% yield reduction, respectively, compared to the fully fertilized treatment with manure.

### Correlation of variables

When correlating root and shoot variables (all dates), a positive correlation (0.7) was observed between shoot and root biomass, beside others variables (Figure S6a). The tiller number correlated positively (0.4) with yield, more than all other variables including shoot biomass (0.2). The root biomass in dates 1 to 4 was strongly associated with final yield (0.6-0.9) and moderately on date 5 (0.4) (Figure S6b-S6e). The correlation of root angle and number of nodal roots with yield varied widely in direction and magnitude of the correlations from date to date. The number of nodal roots was correlated positively with yield on dates 1 and 2 but negatively on dates 3 and 4 (all dates: 0.3). This led to a week positive correlation of root biomass (0.3) and root angle (0.2) with final grain yield for all sampling dates.

## 5) Discussion

Our study presented the effects of N and P fertilizer omission on winter rye shoot and root growth from tillering to flowering. The N and P omission led to a decrease in shoot biomass over time, with stronger reduction in the N omission treatment particularly in the last two sampling dates. Similarly for root biomass, the N omission was more detrimental than the P omission treatment, showing the lowest root biomass among treatments. Also, a decrease of root biomass in the fully fertilized treatment in date 5 was not reflected in shoot biomass, which indicates an alteration in above and below ground allocation of biomass. The R/S ratio was the highest in the N and P omission treatments, which is similar to the findings by Lopez et al. (2023) who reported increased R/S ratio under N stress, nut also reported inconsistent responses in R/S for P stress conditions.

With regards to root morphological traits, a positive but only moderate correlation between the number of nodal roots and grain yield was observed. Also, during the last two sampling dates, the number of nodal roots had a tendency to be higher in the P omission treatment, compared to the fully fertilized treatment with manure. A meta-analysis from (Niu et al., 2013), reported similar findings, where P omission promoted lateral root growth in cereal crops. Grando and Ceccarelli (1995), also compared modern barley cultivars, landraces and wild barley (*Hordeum spontaneum* L.) and showed that there was a significant increase in the number of seminal roots during domestication, suggesting that there may be a relationship between seminal root number and crop productivity.

P omission tends to decrease the tiller number (Graham et al., 1983; Rodríguez et al., 1999). In our results, a trend to decreased tiller number in N and P omission treatments compared to the NPKCa+m+s treatment was observed, though not-significant for dates 1 and 2, but significant for date 3. However, in date 5, the opposite trend was observed where the P omission treatment resulted in higher tiller numbers compared to the NPKca+m+s treatment. In general, under P omission, roots tend to have a wider root angle, due to shallower and broader roots (Bonser et al., 1996; Niu et al., 2013). This trend was not as clear in our results, as, when significant differences were found, P omission treatment showed a non-significant difference when compared to the fully fertilized treatment. Robinson et al. (2018) reported that root angle was more strongly associated with yield than root number. This was the case in our study for sampling dates 3 and 5 but not in the rest of sampling dates. N omission also showed more steep root angles around flowering (sampling date 5), this shift suggests a strategy to explore more soil volume and thus, allow more resource acquisition (Trachsel et al., 2013). The effect of N fertilizer on spring barley root phenotypes was assessed by Siddiqui et al. (2021) showing a tendency for longer and narrower root angles under manure application in an organic system, compared to the lines grown with mineral fertilizer N supply, which showed shorter and wider root system. In line with that we found that N omission treatment showed more steep root angles around flowering.

In plant breeding programs, root traits are generally less considered for selection because root traits carry low heritability, are variable in expression across environments, and require labor-intensive field measurements (Purushothaman et al., 2017). However, some studies state that architectural traits like root preference for shallow or deep soil layers, root angle, and lateral branching are under strong genetic control (El Hassouni et al., 2018; Lynch, 2007). El Hassouni et al. (2018) tested 25 durum genotypes at five locations with different water regimes. All traits connected to root angle showed a very high heritability and were not affected by the water scarcity after anthesis. In contrast, Robinson et al. (2018) tested 216 spring barley breeding lines in pots and found a genetic relationship between seminal root traits and yield (including field data from 20 sites), but the direction and magnitude of the correlations varied across the environments. In our study, we found a high variability of the root traits across dates and the treatment ranking per date. A rather strong positive correlation was observed between shoot and root biomass, as well as root biomass with final yield. In declining order, the number of tillers, root biomass, and root angle were associated with final yield across sampling dates.

We **conclude** that the environment and development stage at sampling have a strong impact on winter rye root phenotypic plasticity and fertilization. Effects of fertilizer omission on the shoot are not only easier to determine but also clearer in terms of direction and treatment ranking. However, roots play a critical role in plant adaptation to abiotic stresses, with root characteristics being central to soil exploration and nutrient acquisition. Strategic adjustments in root growth patterns e.g. via breeding or improved site-specific cultivar selection are needed to enable plants to optimize resource use efficiency in suboptimal environments.

## Funding

The presented study has been funded by the Federal Ministry of Food and Agriculture (BMEL) based on a decision of the Parliament of the Federal Republic of Germany via the Federal Office for Agriculture and Food (BLE) (grant number 2822ABS010), the Deutsche Forschungsgemeinschaft (DFG, German Research Foundation) under Germany’s Excellence Strategy-EXC 2070-390732324 (PhenoRob), by DFG – SFB 1502/1 – 2022 project number: 450058266, the Federal Ministry of Education and Research (BMBF) (project “Sustainable Subsoil Management-Soil3, Grant 031B0151A), as well as by the European Union (EU horizon project IntercropVALUES, grant agreement No 101081973).

## Notes

### Competing Interest Statement

The authors have declared no competing interest.

