## supplement for "Winter rye root growth and plasticity in response to nitrogen and phosphorus omission under field conditions": Supplementary material_WinterRyeNPomission.pdf

Journal: Frontiers in Plant Sc
Brief research report

**Winter rye root growth and plasticity in response to nitrogen and phosphorus omission under field conditions.** ***By Nyameasem et al.***

**Supplementary Material**

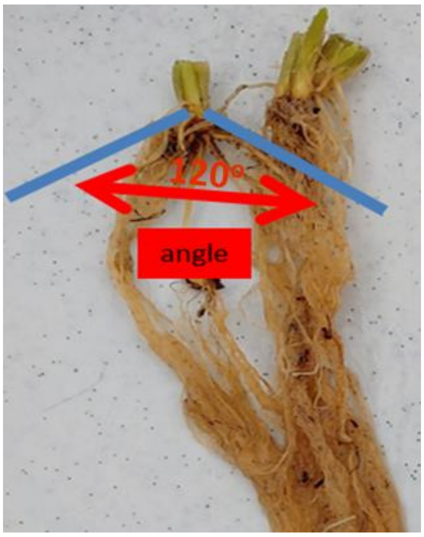


**Figure S1.** Depiction of root angle measurements (blue line) in rye plants, root angle is measured in the nodal roots.

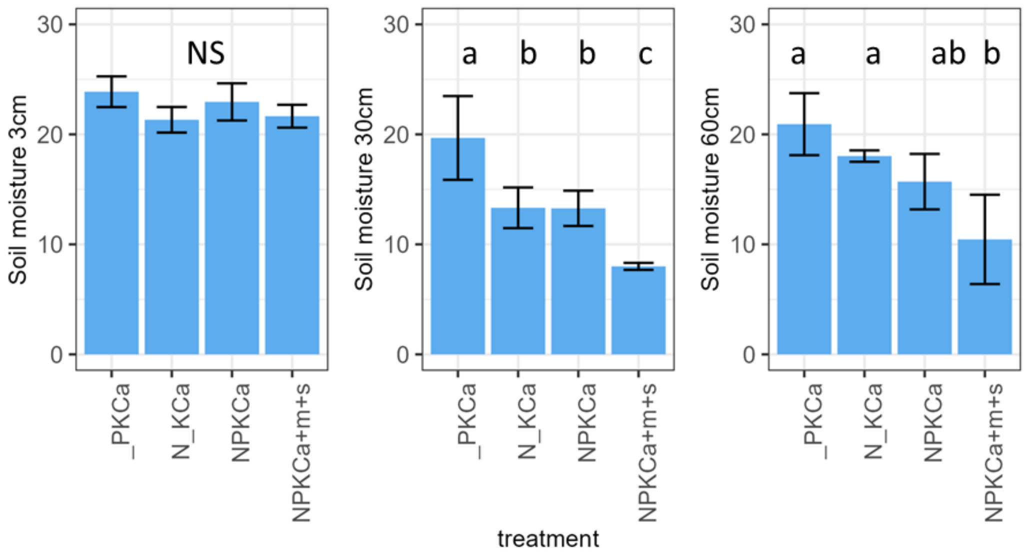


**Figure S2.** Soil moisture content (%) at 3, 30, and 60 cm soil depth on 27/05/2022, around flowering as affected by nitrogen and phosphorus treatments during the 2022 season, over five sampling dates (1: 16/03/2022, 2: 04/04/2022, 3: 29/04/2022,4: 27/5/2022 and 21/06/2022). Treatments: Fully fertilized plus manure (NPKCa+m+s), fully fertilized with mineral fertilizer only (NPKCa), N omission (_PKCa) and P omission (N_KCa). Values followed by the same letter do not differ according to Tukey high significant difference at 5% confidence level. NS= not significant at 5% confidence level.

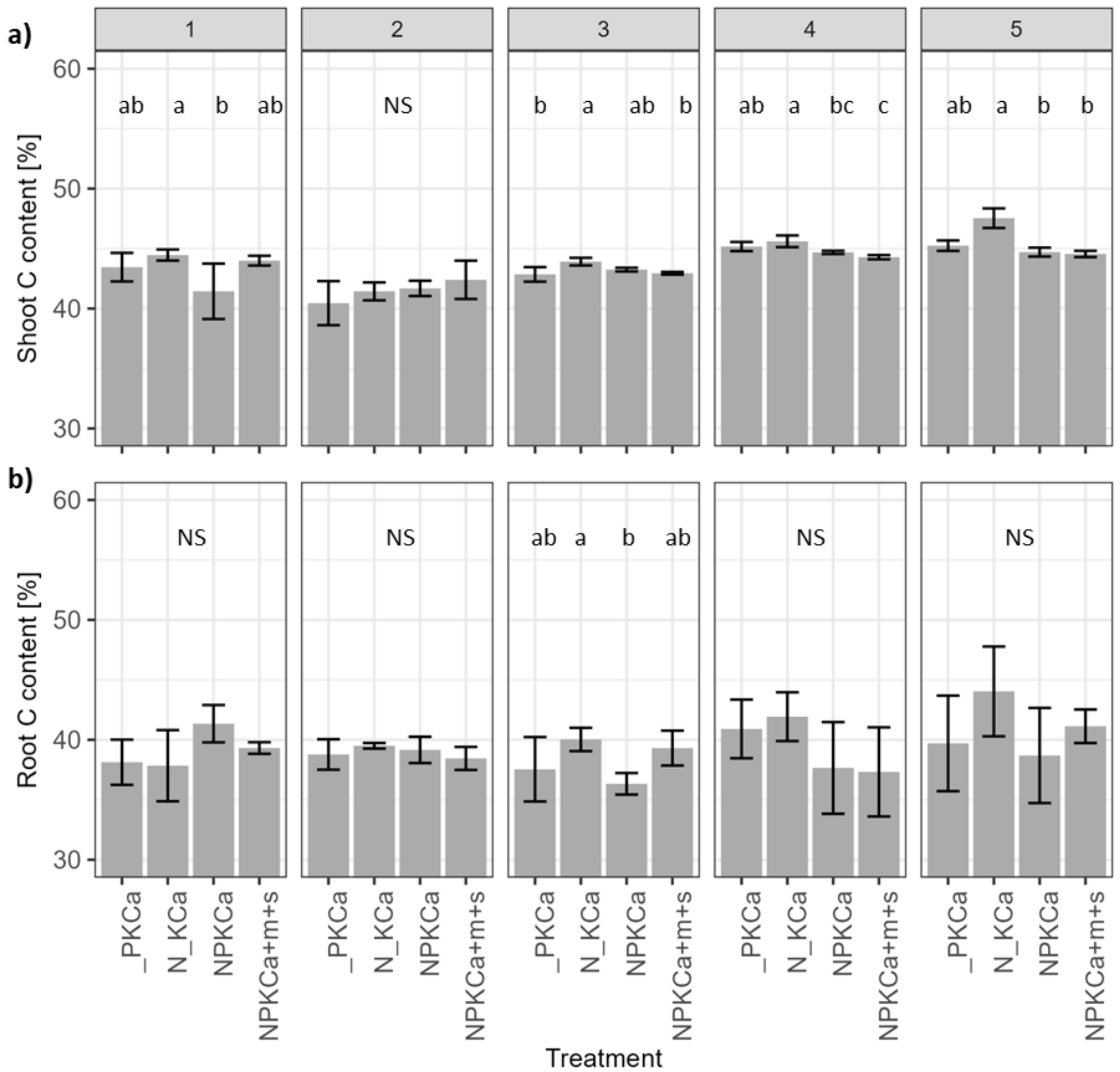


**Figure S3.** Shoot (a) and root (b) C content (%) as affected by nitrogen and phosphorus treatments during the 2022 season, over five sampling dates (1: 16/03/2022, 2: 04/04/2022, 3: 29/04/2022,4: 27/5/2022 and 21/06/2022). Treatments: Fully fertilized plus manure (NPKCa+m+s), fully fertilized with mineral fertilizer only (NPKCa), N omission (_PKCa) and P omission (N_KCa). Values followed by the same letter do not differ according to Tukey high significant difference at 5% confidence level. NS= not significant at 5% confidence level. For shoot N content in date 5, an Aligned rank transform for nonparametric factorial ANOVA was implemented.

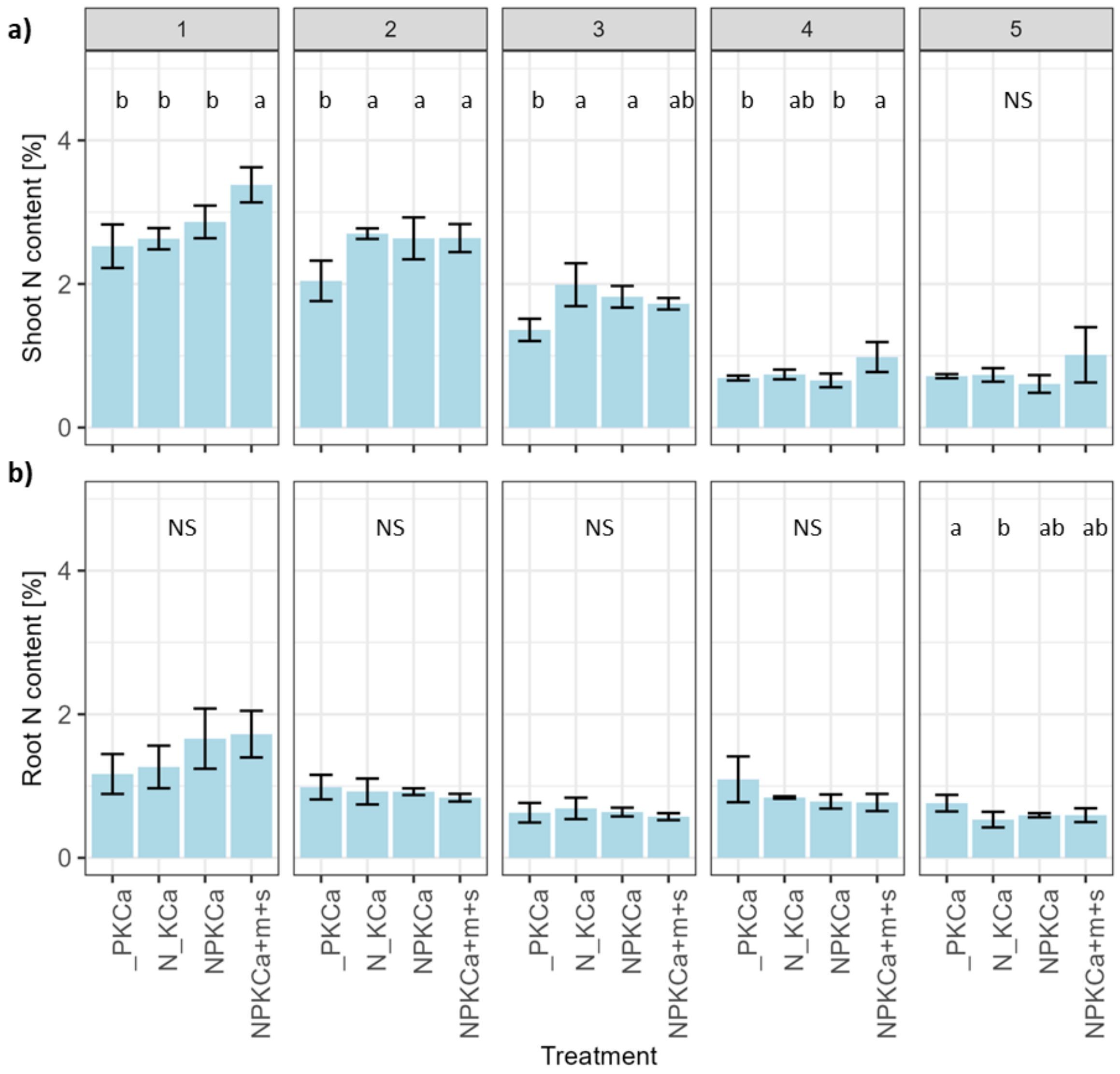


**Figure S4.** Shoot (a) and root (b) N content (%) as affected by nitrogen and phosphorus treatments during the 2022 season, over five sampling dates (1: 16/03/2022, 2: 04/04/2022, 3: 29/04/2022,4: 27/5/2022 and 21/06/2022). Treatments: Fully fertilized plus manure (NPKCa+m+s), fully fertilized with mineral fertilizer only (NPKCa), N omission (_PKCa) and P omission (N_KCa). Values followed by the same letter do not differ according to Tukey high significant difference at 5% confidence level. NS= not significant at 5% confidence level. For root N content in date 4, an Aligned rank transform for nonparametric factorial ANOVA was implemented.

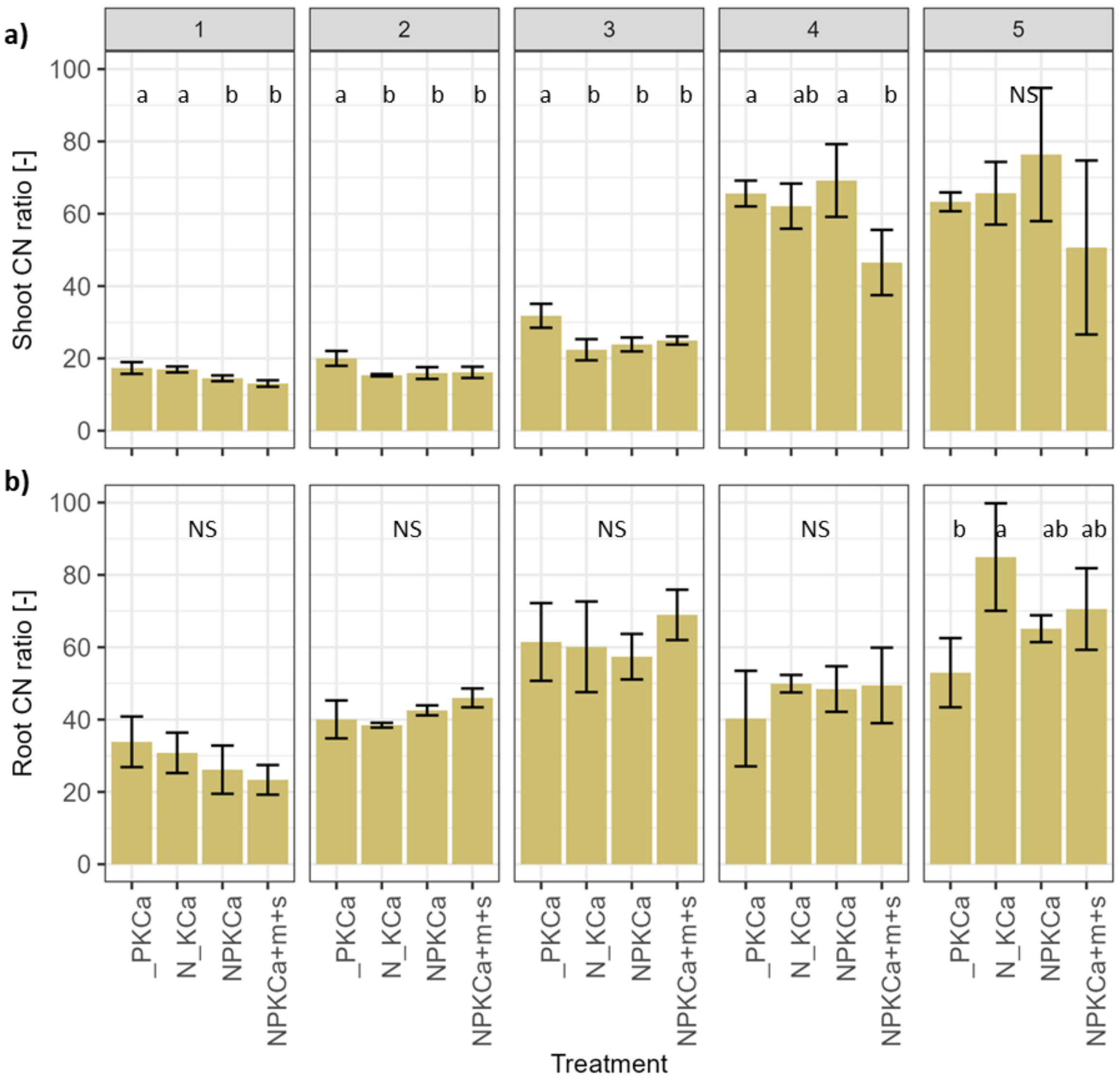


**Figure S5.** Shoot (a) and root (b) C:N ratio as affected by nitrogen and phosphorus treatments during the 2022 season, over five sampling dates (1: 16/03/2022, 2: 04/04/2022, 3: 29/04/2022,4: 27/5/2022 and 21/06/2022). Treatments: Fully fertilized plus manure (NPKCa+m+s), fully fertilized with mineral fertilizer only (NPKCa), N omission (_PKCa) and P omission (N_KCa). Values followed by the same letter do not differ according to Tukey high significant difference at 5% confidence level. NS= not significant at 5% confidence level.

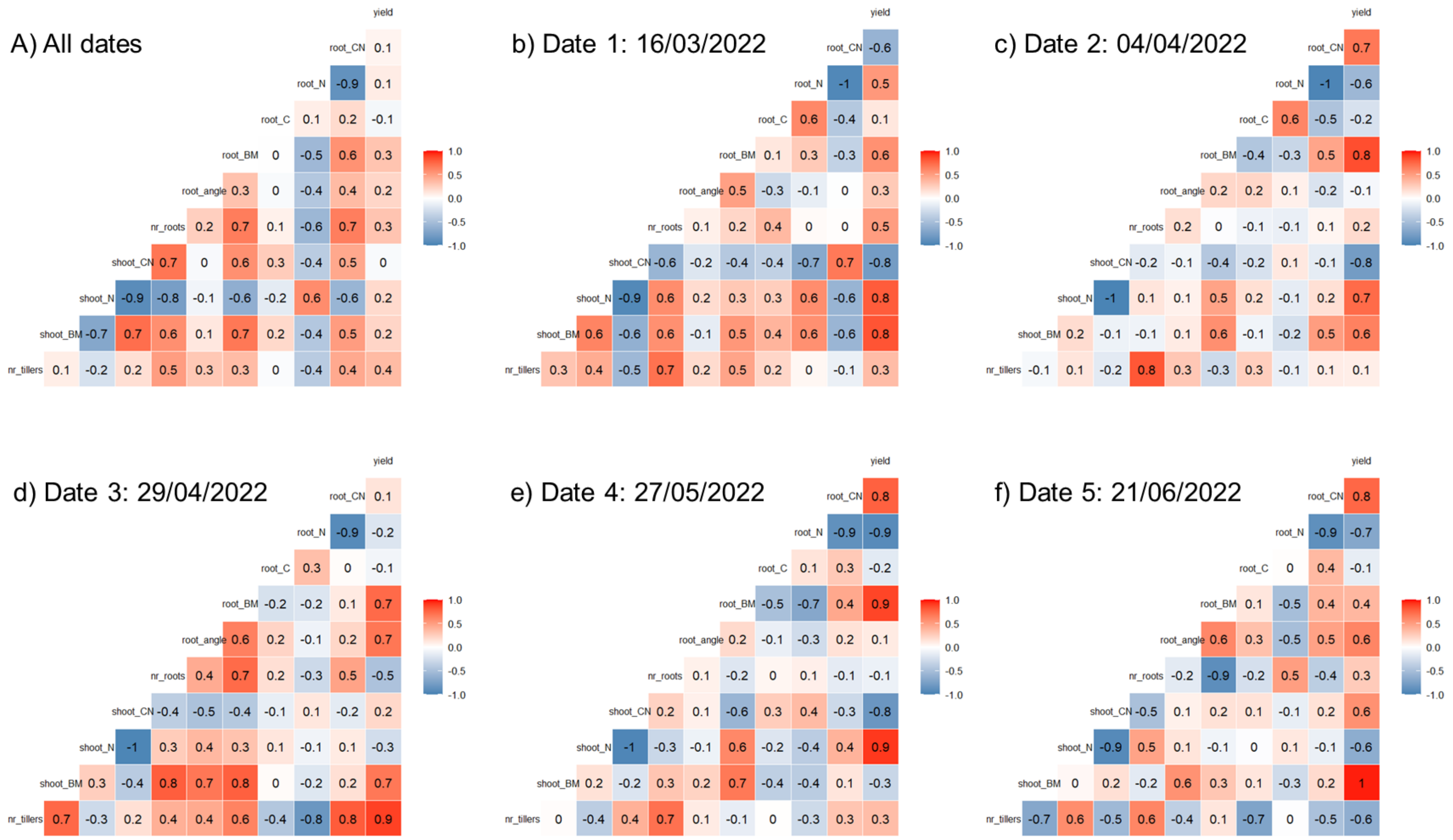


**Fig. S6:** Correlation coefficients for different root and shoot variables for a) all five sampling dates (1: 16/03/2022, 2: 04/04/2022, 3: 29/04/2022,4: 27/05/2022 and 21/06/2022) and, b to f) by sampling date. Abbreviations= nr_tillers= number of tillers (no.), shoot_BM= shoot biomass (g $m^{-2}$), shoot_N= shoot nitrogen content (%), shootCN= Shoot C:N ratio (-), nr_roots= number of seminal roots (no.), root_angle= root angle (no.), root_BM= root biomass (g $m^{-2}$), root_C= root C content (%), root_N= root N content (%), root_CN (%), yield (g $m^{-2}$).
